# A New Technique and Device for Controlled and Continuous Drug Delivery into the Brain – A Proof of Concept Study

**DOI:** 10.1101/746966

**Authors:** U.R. Anoop, Kavita Verma

**Author notes:** **Correspondence to:** Dr. Anoop.U.R, MDS, UR Anoop Group, Pondicherry, India. 605008.

## Abstract

**Background:** Drug delivery into the brain has been a challenge for the past 100 years because of the blood brain barrier. The existing non-invasive techniques cannot provide controlled and continuous drug delivery into the brain and the invasive techniques make the brain prone to infection from external agents. Hence a new technique which can provide controlled and continuous drug delivery without the need for any surgical intervention in the brain holds immense potential.

**Objective:** The objective of this study is to deliver drugs into the brain using a novel oral and maxillofacial technique and device.

**Method:** Drug delivery into the brain from the oral and maxillofacial region was tested using a novel technique and device in an in vivo rabbit model and an ex vivo goat head model. A control animal and an experimental animal were used in each study. Drugs which do not cross the blood brain barrier normally were tested. Dopamine was delivered in vivo from the maxillo-facial region. Anti-glial fibrillary acidic protein antibody was delivered ex vivo from the oral region. Samples were collected from different sites including the brain and the optic nerve.

**Results:** The in vivo model showed a significant increase of dopamine at the pons (51.89%), midbrain (27%), medulla (48.5%) and cortex (72.637%). On including samples from other regions in the t-test, the increase was not statistically significant (*p*=0.538), suggestive of a central feedback mechanism for brain and peripheral dopamine. A decrease in plasma dopamine during drug delivery further supported a central control for dopamine. In the ex vivo model, a statistically significant (*p*=0.047) delivery of antibodies occurred at multiple sites including pons (86.7%), cortex (256.5%), and the optic nerve (128.8%).

**Conclusion:** This technique and device can deliver drugs into the brain without detectable increase in systemic circulation. Therefore it may be used for delivering drugs in Parkinson’s disease, Alzheimer’s disease, Pain management, Brain tumors especially pontine tumors, infections like neuro-AIDS, Basal meningitis etc. Retinal drug delivery may also be possible.

## BACKGROUND

It is more than 100 years since the blood brain barrier was discovered [1]. The blood brain barrier protects the brain from external agents. But this natural protection also prevents 98% of the small drugs and 100% of the large drugs available today from entering the brain in concentrations that can produce the required therapeutic effects [2].

In Parkinson’s disease, focus is currently on techniques to effect continuous tonic firing of diseased dopamine neurons as in normal physiologic conditions. But dopamine does not cross the blood brain barrier normally.

In Alzheimer’s disease, focus is currently on immunotherapy. But, the entry of the peripherally administered antibodies into the brain has been limited to around 0.1% to 0.2% by the blood brain barrier.

In tumors involving the brainstem, the prognosis has been poor because drug delivery is restricted by the blood brain barrier.

In the eye, retinal drug delivery is a challenge till date because of the blood retinal barrier.

This novel device and technique address the issue of delivering drugs into the brain.

The device is implanted in the oral or maxillofacial region and the drugs are delivered directly into the connective tissue of the respiratory mucosa of the maxillary sinus, from where the drugs get transferred into the brain including the optic nerve through the neural, lymphatic and vascular pathways.

This technique is advantageous because it avoids the iatrogenic brain damage caused by the existing invasive techniques like convection enhanced drug delivery. It also provides continuous and controlled drug delivery which is not feasible with the existing non-invasive techniques like nasal drug delivery, receptor mediated transport, ultrasound mediated opening of blood brain barrier etc.

This technique provides access to the optic nerve and vessels and therefore can deliver drugs into the retina and the vitreous.

## MATERIALS AND METHODS

The in vivo proof of concept study was done using a rabbit model in an OECD GLP certified pre-clinical research lab with a CPCSEA approved animal lab facility and accredited by the Association for Assessment and Accreditation for Laboratory Animal Care International. The experimental protocols were approved by the Institutional Animal Ethics Committee of the pre-clinical research lab. All methods were performed in accordance with the relevant guidelines and regulations.

### In Vivo animal model

Two New Zealand Male White Rabbits aged 4 to 6 months and weighing around 2 Kg were used as the proof of concept models to study the delivery of dopamine into the brain using the new device and technique. The experimental model and the control model were housed individually in stainless steel wire mesh cages under a 12 hour light/dark cycle. Food and water was provided ad libitum to the animals.

### The Technique

The rabbit models were anaesthetized using a combination of xylazine (5 mg/kg bwt/Intramuscular) and ketamine (35 mg/kg bwt/ Intramuscular). Following induction of anesthesia, the animals were placed on a flat surface in the prone position.

A midline incision was made on the nasal dorsum. The skin and periosteum were lifted to expose the nasal bone and nasoincisal suture line. A round bone window of about 0.5 cm was made over the maxillary sinus region using a bone trephine drill unilaterally. The overlying bone was removed without tearing the underlying respiratory mucosa. The respiratory mucosa was then gently released from the bony margins and the prototype of the device was placed into the bone window. The other nasal cavity was left patent for airway maintenance. The anaesthetized rabbits were monitored carefully during experimental time for anesthetic perception by pedal/pinna/corneal reflex.

Figure 1 shows the experimental model where dopamine was given in a controlled and continuous manner using an external drug infusion pump for 20 minutes at the rate of 23 microliter/minute. Around 1.0 ml of blood was collected from the ear vein at 0, 10, 20 and 30 minutes into tubes containing 100 IU/ml Heparin sodium. Plasma was isolated by centrifuging at 3000 rpm for 5 minutes. The isolated plasma was stored at −80°C.

**Figure 1.**
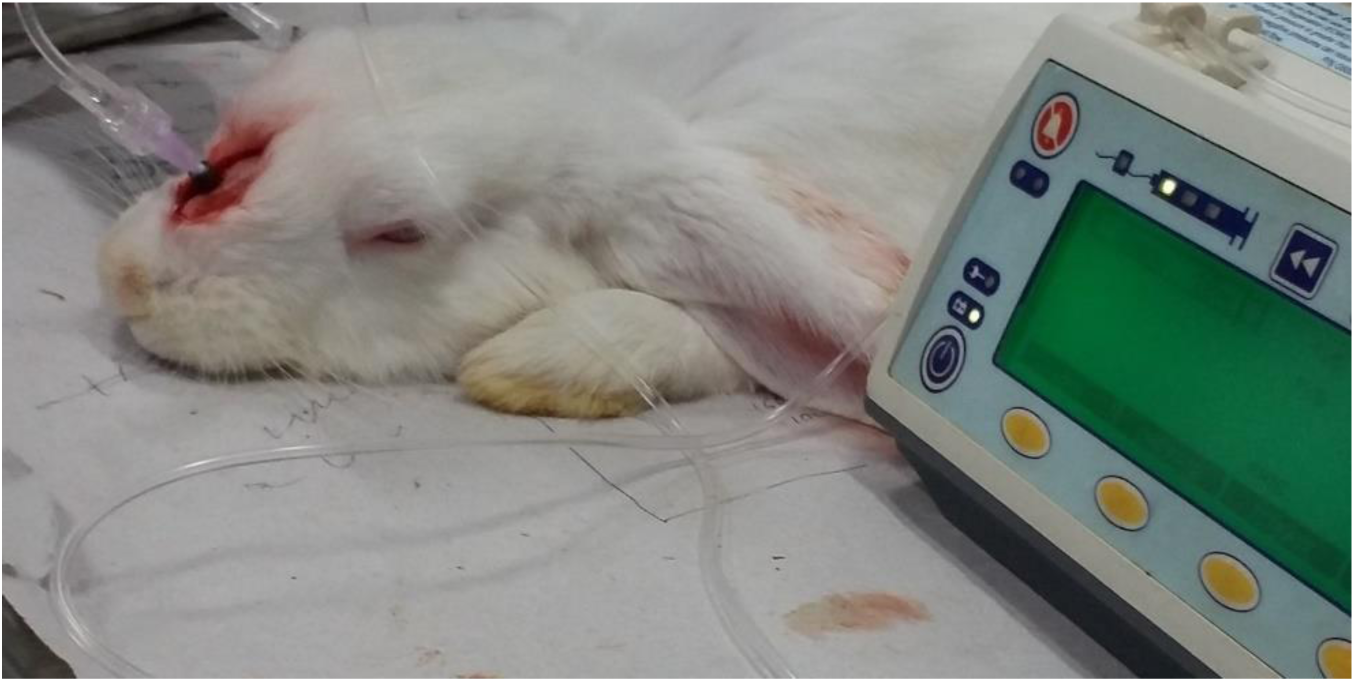
shows the experimental model where dopamine is given in a controlled and continuous manner using an external drug infusion pump.

In the control model, anesthesia was given and the bone window was prepared. But, no dopamine was infused. It was used for studying the normal dopamine concentration in the brain tissue.

Cardiac perfusion was done using saline to clear the brain of any residual blood under deep surgical anesthesia. Before and during the perfusion, the animal was monitored visually for withdrawal reflex. The perfusion was carried out in necropsy hood using a peristaltic pump to deliver the warm physiological saline at controlled speed. The left ventricle was identified and a small cut was given at the apex to insert the perfusion cannula into the ventricle till it reached the aorta. The inserted cannula was then clamped in place and normal saline was infused. The right atrium was then opened and the animal was perfused with normal saline for 15 to 20 minutes to remove the blood completely.

Once, most of the visceral organs turned pale in color, perfusion was stopped. The cleared brain was removed carefully and stored in dry ice. Duramater adhering to the cranial cavity and the nasal mucosa were removed and stored in dry ice. The tissue samples were homogenized with buffer, centrifuged and subjected to HPLC. The plasma samples were also subjected to HPLC.

### Ex-Vivo animal model

The blood brain barrier has been found to be structurally maintained after death for a few hours at least in relation to large molecules [3]. Therefore, it was decided to use a goat head which was procured immediately after decapitation from an abattoir to assess the feasibility of delivery of large antibodies across the blood brain barrier using the new technique and device. An experimental model and a control was used. The goat heads were collected immediately after decapitation and transferred to the lab within 20 minutes of decapitation. In the study model, the surgical process was completed within 10 minutes and 2.5 ml of polyclonal rabbit anti glial fibrillary acidic protein antibody from DAKO (Lot:20044594) was delivered slowly under the connective tissue of the respiratory mucosa of the palatine sinus using the device, over a period of 20 minutes.

### The Technique

Figure 2 shows the intra-oral surgical technique. A crevicular incision was placed along the cervical margins of the posterior maxillary teeth on the palatal aspect and a muco-periosteal flap was elevated to expose the palate. A bone window was created on the bone overlying the palatine sinus to expose the connective tissue of the palatine respiratory mucosa. The mucosa was gently released from the surrounding bone margins. The prototype of the device was placed into the surgical site without tearing the mucosa. 2.5 ml of polyclonal rabbit anti glial fibrillary acidic protein antibody was injected slowly using a syringe over a period of 20 minutes.

**Figure 2.**
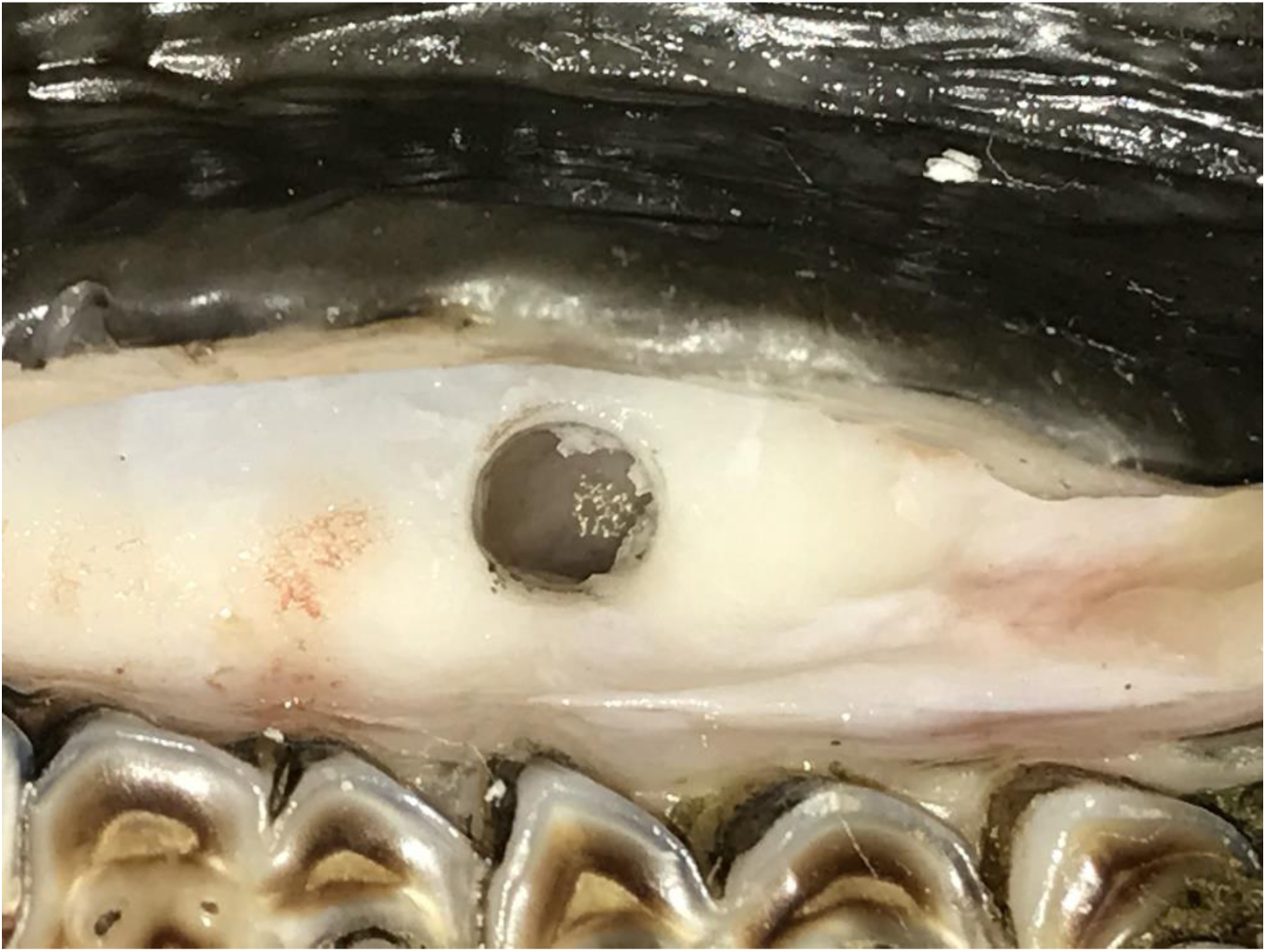
shows the intra-oral technique.

The brain was removed within 20 minutes and stored in dry ice. Duramater adhering to the cranial cavity was removed and stored in dry ice. Similarly the nasal mucosa adjacent to the implantation site of the device was resected and stored in dry ice. The eye from the same side was enucleated along with part of the optic nerve and stored in dry ice. Vitreous was aspirated from the eye using a 5ml syringe through a trans-scleral approach and stored in dry ice. The tissue samples were homogenized with buffer, centrifuged and subjected to HPLC.

## RESULTS

### IN VIVO RABBIT MODEL-DOPAMINE

Figure 3 compares the concentration of dopamine in the brain, duramater and the respiratory mucosa of the maxillary sinus between the experimental model and the control. Dopamine concentration in Midbrain, which is the target for Parkinson, showed an increase of 27% when compared with the control. Dopamine was also increased in Cortex (72.637%), Cerebellum (21.75%), Medulla (48.5%) and Pons (51.8%) when compared with the control. A significant finding was that the Diencephalon showed a decrease of −27.76% when compared with the control. Moreover the respiratory sinus mucosa also showed a decrease of −2.454%. The Olfactory Bulb and Tract showed a decrease of −6.1238% and the Optic Nerve showed a decrease of −15.746%.

**Figure 3.**
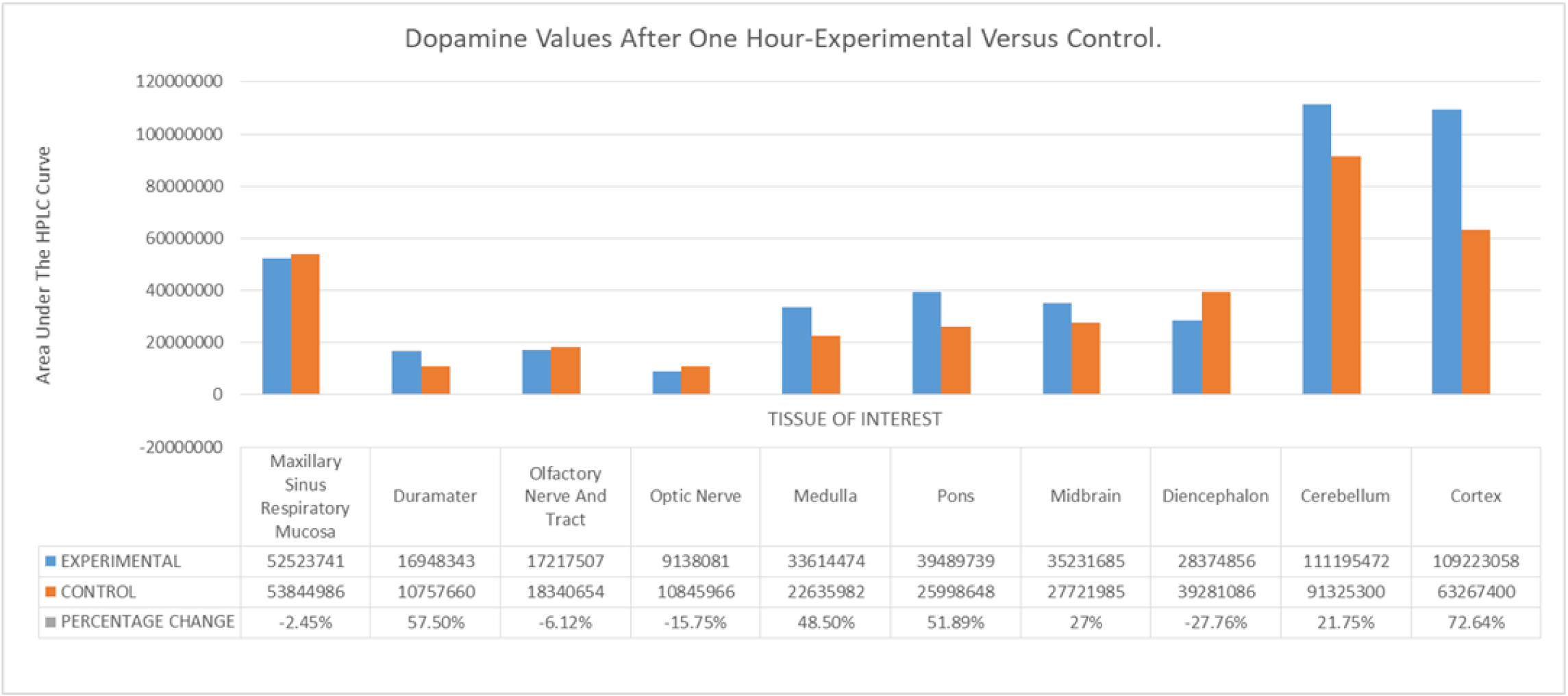
compares the concentration of dopamine in the brain, duramater and the maxillary sinus respiratory mucosa between the experimental model and the control.

Inspite of a significant increase in dopamine in the brainstem and the cortex, a t –test performed using SPSS software version 20.0 (SPSS Inc., Chicago, IL) found the changes to be not statistically significant (*p*=0.538). This finding suggests that a central feedback control for brain and peripheral dopamine may be present.

Figure 4 compares the concentration of dopamine in the blood in the experimental model during the drug infusion period. The blood values showed a decrease in dopamine levels at 10 minutes and 20 minutes interval which correlated with the time the drug was being infused. Once the dopamine infusion was stopped at 20 minutes, the blood values again peaked at 30 minutes. The peak was higher than that at 0 minutes

**Figure 4.**
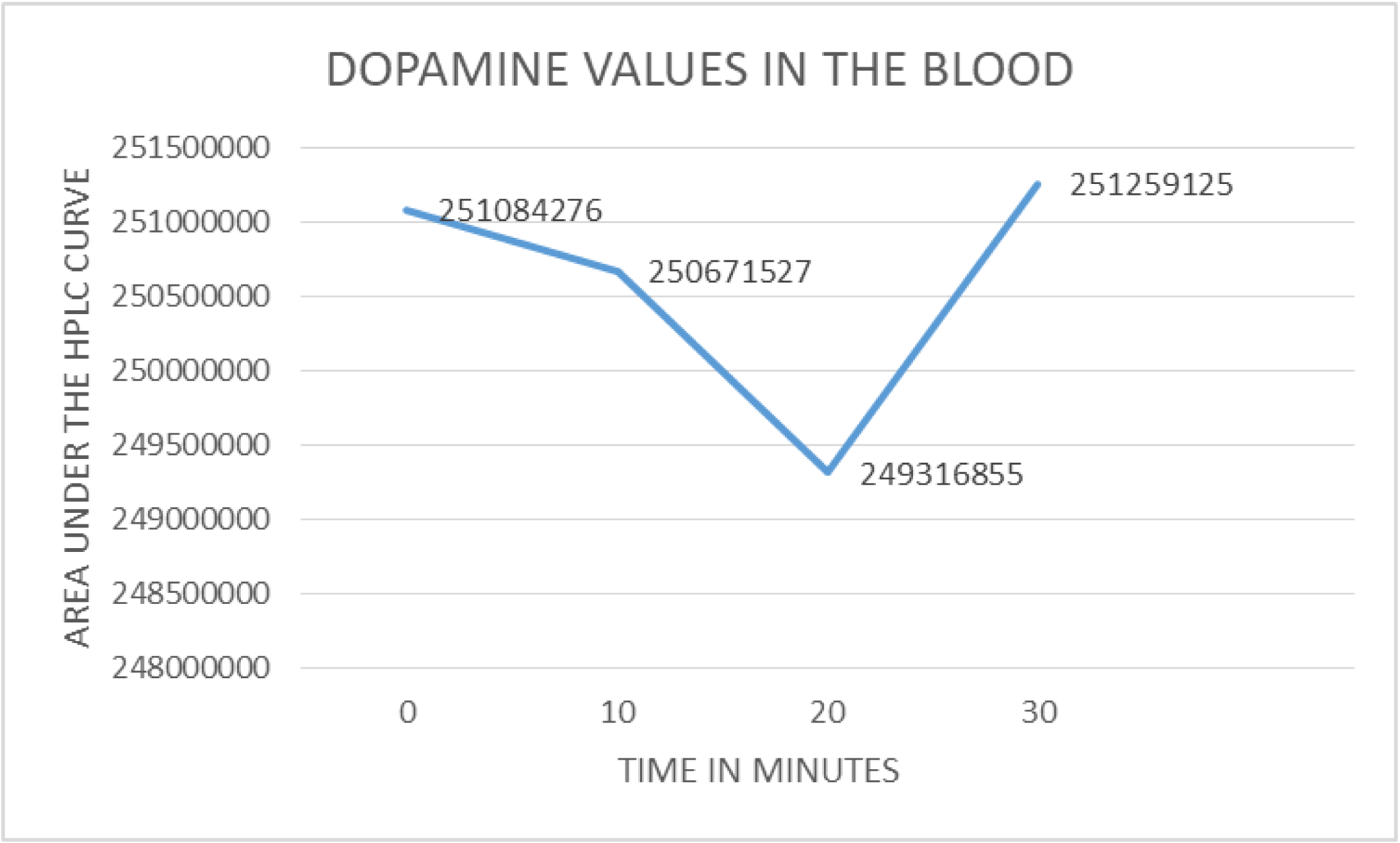
compares the concentration of dopamine in the blood in the experimental model during the dopamine infusion period.

### EX VIVO GOAT HEAD MODEL-POLYCLONAL RABBIT ANTI-GFAP ANTIBODY

Figure 5 compares the concentration of anti-gfap antibody in the brain duramater, vitreous and the respiratory mucosa of the palatal sinus between the experimental model and the control. In the experimental model, the respiratory mucosa of the palatine sinus showed an increase of 36.97% when compared with the control. The Duramater showed an increase of 22.76%. The Pons showed an increase of 86.7%. The Cortex showed an increase of 256.5%. The Optic nerve showed an increase of 128.8% and the Vitreous showed an increase of 289.67%. A t-test performed using SPSS software version 20.0 (SPSS Inc., Chicago, IL) found the changes to be statistically significant (*p*=0.047)

**Figure 5.**
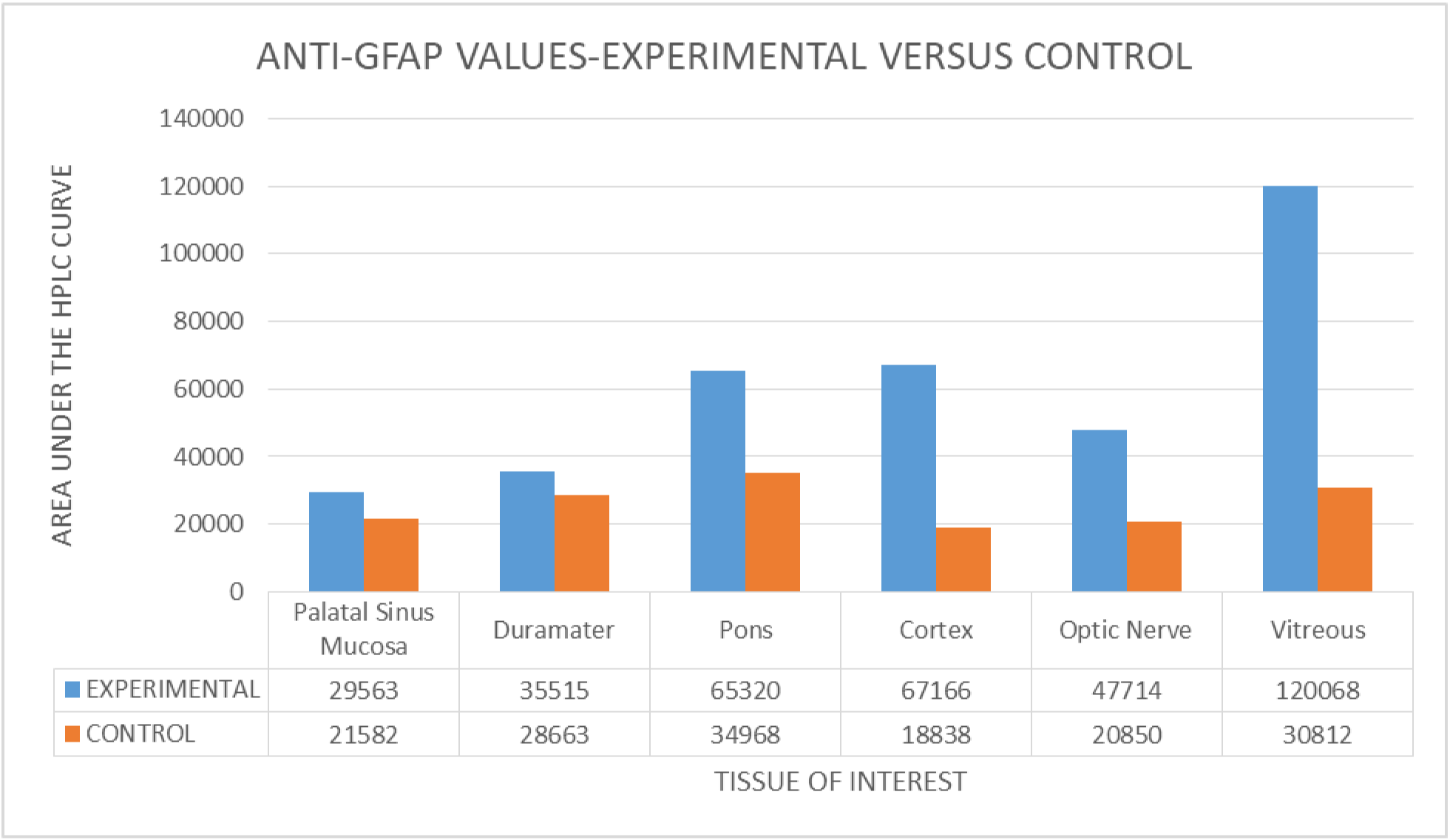
compares the concentration of anti-gfap antibody in the brain, duramater, vitreous and palatine sinus respiratory mucosa between the experimental model and the control.

## DISCUSSION

A proof of concept study was conducted to test the efficacy of a new technique and device for delivering drugs into the brain including the optic nerve. An In-vivo study using a rabbit model and an Ex-vivo study using a freshly decapitated goat head was done. Drugs which normally do not cross the blood brain barrier were selected for the study. Dopamine, a small molecule and Anti-gfap, a large antibody were tested using this device to assess whether drugs of different sizes can be delivered using this technique.

In the in vivo study, the connective tissue underlying the respiratory mucosa of the maxillary sinus was surgically accessed by creating a window on the overlying bone [4]. The prototype of the device was strategically placed with its delivery tip located beneath the respiratory mucosa. A pump was used to deliver dopamine in a controlled and continuous manner.

Trigeminal nerve supplies the nasal respiratory mucosa and the maxillary sinus mucosa and relays at the Pons [5]. Cerebrospinal fluid drains through nasal lymphatics which are free of valves [6]. The veins from parts of the maxillary sinus and the nasal region drain into the pterygoid venous plexus and reach the cavernous sinus [7]; [8]. Studies have shown retrograde dopamine transfer in the cavernous sinus because of counter current mechanism in perfused ex vivo rabbit models [9]. Hence the drug delivered directly into the connective tissue of the respiratory mucosa, may be transported from the nasal and maxillary sinus region through the neural [5]; lymphatic [6]; and the paravascular [10] routes into the brain.

Dopamine concentration in Midbrain, which is the target for Parkinson, showed an increase of 27% when compared with the control. The Cortex showed an increase of 72.637% when compared with control. These values show that these therapeutic sites can be accessed and dopamine can be delivered across the blood brain barrier without any direct surgical intervention in the brain. Moreover sufficient concentration of the drug can be maintained at the site for producing the therapeutic effect. The significant increase shows that very low concentration of drugs may be used to produce therapeutic effects in the brain. By using controlled drug delivery or slow release formulations it will be possible to provide continuous drug delivery into the brain for a longer period of time. This technique can therefore produce continuous tonic stimulation of the dopamine receptors. Duramater showed an increase of 36.52%. The infused dopamine may be delivered by retrograde paravascular transport through branches of the maxillary artery and through the pterygoid venous plexus into the duramater. Moreover Duramater is also supplied by afferent branches from the trigeminal ganglion and the sphenopalatine ganglion in the anterior and middle cranial fossa. Therefore neural pathway also may play a role in the delivery of the infused dopamine. Lymphatic pathway also may be involved.

Cerebellum showed an increase of 21.75%. The Medulla and Pons showed an increase of 48.5% and 51.8% respectively. The Trigeminal enters the brainstem at the Pons. Therefore the structures adjacent to the pons also may also receive the infused drug. Moreover it shows that delivery of the drug occurs in the rostral as well as the caudal part of the brainstem. This may of significance in the treatment of Pontine tumors.

The concentration of dopamine in different brain regions of the control model decreased in the following order, Cerebellum > Cortex > Diencephalon > Midbrain > Pons > Medulla > Olfactory Bulb and Tract > Optic Nerve. But the concentration of dopamine in different brain regions of the experimental model decreased in the following order, Cerebellum > Cortex > Pons > Midbrain> Medulla >Diencephalon > Olfactory Bulb and Tract > Optic nerve. This finding shows that the brainstem of the study model has increased dopamine especially at the Pons. This further correlates with the fact that the Trigeminal nerve enters the brainstem at the Pons and therefore tends to deliver more drug at the Pons and the adjacent sites.

A significant finding was that the Diencephalon showed a decrease of −27.76% when compared with the control. This is significant as this decrease could be a result of the feedback from the cortical, subcortical and brainstem regions which received the infused dopamine in excess. This further signifies the opportunity to use lower concentration of the drug to produce therapeutic effects in the brain.

Moreover the dopamine concentration in the respiratory sinus mucosa showed a decrease of −2.454% when compared with the control. This finding is significant because even when dopamine was delivered directly to the connective tissue of the respiratory mucosa in the experimental model, the net result was a decrease in dopamine. Studies have shown that there is minimal metabolism of dopamine in the nasal mucosa and trigeminal pathway is an effective mode of transport [11]. The respiratory mucosa is predominantly supplied by the branches of the trigeminal nerve. D1 and D2 receptors are seen in the trigeminal ganglion and spinal trigeminal nucleus. Studies have shown that dopamine has an inhibitory effect on the trigeminal nucleus. Moreover centrally active D2 antagonist had a stimulatory effect and reversed the effect of dopamine, while peripherally active D2 antagonist had no effect on the trigeminal nucleus [12]. Therefore the reason for the decrease in respiratory mucosal dopamine may be a feedback due to the delivery of excess dopamine in the brainstem region.

Similarly, the Olfactory Bulb and Tract showed a decrease of −6.1238% when compared with the control. The increase in dopamine in the primary olfactory relay area in the cortex or decrease in the secondary relay area in the thalamus may have caused a decrease in the Olfactory Bulb and Tract dopamine by a feedback mechanism. This finding may be significant in analyzing the cause and treatment options for olfactory dysfunction in Parkinson’s [13].

The Optic Nerve showed a decrease of −15.746% when compared with the control. The retina and the optic nerve have dopaminergic innervation. The optic pathway from the retina relays in the lateral geniculate ganglion in the Thalamus which further relays in the visual cortex. The retinal efferent also relays in the pulvinar of the thalamus, superior colliculus, and midbrain tectum. Therefore excess dopamine in these sites can decrease dopamine in optic nerve by a feedback mechanism. In this study, the increase in dopamine in the midbrain, the cortical and the sub-cortical sites along with the decrease in dopamine in the diencephalon may have caused a decrease in the optic nerve dopamine by a feedback mechanism. Visual dysfunction is also seen in Parkinson’s [14]; [15]. Hence this finding may be significant.

In spite of significant changes at multiple sites, the dopamine variation in the experimental model was not statistically significant when compared with the control. This further emphasizes the innate tight control by the feedback mechanism of the brain. The feedback mechanism may maintain the total brain dopamine within a range.

Similarly, the blood values showed a decrease of −0.16% in dopamine levels at 10 minutes. The dopamine values dipped further to −0.70% at 20 minutes. It correlated with the time the drug was being infused (Fig: 4). Once the dopamine infusion was stopped at 20 minutes, the blood values again peaked at 30 minutes. The dopamine value at 30 minutes was higher by 0.069% than that at 0 minutes. This finding is significant because there may be a central control of peripheral dopamine release and peripheral dopamine may be mildly inhibited during the infusion. Moreover this technique does not cause any significant increase of dopamine in the peripheral circulation which may have clinical significance.

It is accepted that the blood brain barrier is maintained at least in relation to large molecules after death. The study by Hossmann and Olson on a cat model found that intravenously injected Evans blue and horse radish peroxidase did not cross the blood brain barrier for up to 3 hours even when acute complete cerebral ischemia was caused by arterial ligation [3]. It has also been noted that the brain is less subject to postmortem redistribution of drugs when compared with other organs [16].

Therefore an ex-vivo study using a goat head following immediate decapitation was done to assess the efficacy of the new drug delivery device and technique. Anti-gfap antibody was delivered under the connective tissue of the palatine respiratory mucosa of a freshly decapitate goat head using the device.

Recently paravascular transport pathways which are analogous to the glymphatic pathways in the brain have been reported in the retinal vasculature and the optic nerve. It has also been postulated that the paravascular transport pathways may provide drainage from the optic nerve to beneath the internal limiting membrane which forms the boundary between the vitreous and the retina [17]. [18] [19].

It has been shown that fluorescent dextran tracers of 10, 40, 70 and 500kDa sizes, when injected into the CSF at the cisterna magna of mice enter the sub arachnoid space of the orbital optic nerve. While the tracers ranging in size from 10kDa to 40kDa were also seen within the optic nerve, the tracers of 70KDa size had only limited distribution within the optic nerve and the tracers of 500kDa size could not be detected [20].

Intravitreal drug delivery studies have shown that the negatively charged inner limiting membrane has pores of approximately 10nm and does not limit retinal drug delivery of compounds below 100kDa [21].

Therefore drugs delivered beneath the internal limiting membrane of the retina through paravascular transport pathways may also reach the vitreous.

The HPLC study result showed that the concentration of the antibody decreased in the following order Vitreous >Cortex > Optic Nerve > Pons > Palatine sinus mucosa > Duramater. In the study model, the antibody concentration in the respiratory mucosa of the palatine sinus increased by 1.3 fold when compared with the control. In the duramater it increased by 1.2 fold and in the pons by 1.8 fold. The cortex showed a 3.5 fold increase. The optic nerve showed a 2.2 fold increase and the vitreous showed a 3.8 fold increase. The increase of antibody in the vitreous is of clinical significance. The drug could have reached the retina by bypassing the blood retinal barrier through the optic nerve and vessels. It may have further entered the vitreous through the inner limiting membrane. The high concentration of the antibody in the optic nerve therefore shows that a perineural drug delivery route to the retina from the brain is a possibility.

## CONCLUSION

The new drug delivery system and technique can be used for continuous and controlled delivery of drugs into the brain without causing any detectable systemic increase. It can be used for a number of medical conditions like Alzheimer’s, Drug addiction, cancer, Pain management, infections like Neuro-Aids, Basal Meningitis Rare Diseases, Posterior eye segment drug delivery etc. Controlled drug delivery into the brainstem for pontine tumors will be an advantage In the brain, an initial slow paravascular and perivascular clearing of amyloid proteins followed by an active dosing may help in effectively clearing amyloid without amyloid related imaging abnormalities in Alzheimer patients. Maintaining a higher concentration of the drug in the cerebral vasculature continuously, without causing systemic overloading may tire out the efflux machinery and ‘open the gates’ by producing a ‘mob’ effect.

Retina and the central nervous system have a number of similarities that include a common origin of development from the neural tube, presence of a blood-retinal barrier and a blood-brain barrier respectively and the occurrence of age related extra cellular deposits comprising βamyloid referred to as drusen in the retina and as senile plaques in the brain [22]. As it has been shown that the retina and the vitreous can be accessed through the optic nerve, this device and technique can be used for treating diseases of the eye too.

The goat head may be useful as an ex-vivo model for testing delivery of large molecules into the brain along with in vitro models.

As this study is a proof of concept study, further studies with larger sample sizes will be required in the future.

## ACKNOWLEDGEMENT

We acknowledge Dr. D. Saravanan, Phd, co-ordinator, National College Instrumentation Facility, Tiruchirapalli, India and Mrs. S. Kavitha, M Pharm, HPLC analyst, National College Instrumentation Facility, Tiruchirapalli, India for their invaluable support in completing this study.

## AUTHOR’S CONTRIBUTION

Both the authors have contributed equally to the work

## FUNDING

This research did not receive any specific grant from funding agencies in the public, commercial or not for profit sectors.

## COMPETING INTERESTS

The authors, Anoop.U.R and Kavita Verma are stated as the inventors and applicants in the published patent applications US15/177,347 and PCT/IB2016/053899 with National Phase Entry into European Patent Office, Australia, Canada and India. There are no other conflict of interests.

## DATA AVAILABILITY

Data analyzed during the study are included in the article.

## Notes

#### Summary of Updates

Author affiliation Updated

